# A highly resolved integrated single-cell atlas of human breast cancers

**DOI:** 10.1101/2025.03.13.643025

**Authors:** Andrew Chen, Lina Kroehling, Christina S. Ennis, Gerald V. Denis, Stefano Monti

**Affiliations:** Section of Computational Biomedicine, Boston University Chobanian and Avesidian School of Medicine, Boston, MA 02118, USA; Bioinformatics Program, Center for Computing & Data Sciences, Boston University, Boston, MA 022158, USA; Boston University-Boston Medical Center Cancer Center, Boston University Chobanian and Avesidian School of Medicine, Boston, MA, 02118, USA; Section of Hematology and Medical Oncology, Department of Medicine, Boston University Chobanian and Avesidian School of Medicine and Boston Medical Center, Boston, MA, 02118, USA; Department of Pharmacology and Experimental Therapeutics, Boston University Chobanian and Avesidian School of Medicine, Boston, MA, 02118, USA; Department of Biostatistics, Boston University School of Public Health, Boston, MA 02118, USA

## Abstract

In this study, we developed an integrated single cell transcriptomic (scRNAseq) atlas of human breast cancer (BC), the largest resource of its kind, totaling > 600,000 cells across 138 patients. Rigorous integration and annotation of publicly available scRNAseq data enabled a highly resolved characterization of epithelial, immune, and stromal heterogeneity within the tumor microenvironment (TME). Within the immune compartment we were able to characterize heterogeneity of CD4, CD8 T cells and macrophage subpopulations. Within the stromal compartment, subpopulations of endothelial cells (ECs) and cancer associated fibroblasts (CAFs) were resolved. Within the cancer epithelial compartment, we characterized the functional heterogeneity of cells across the axes of stemness, epithelial-mesenchymal plasticity, and canonical cancer pathways. Across all subpopulations observed in the TME, we performed a multi-resolution survival analysis to identify epithelial cell states, and immune and stromal cell types, which conferred a survival advantage in both The Cancer Genome Atlas (TCGA), METABRIC, and SCANB. We also identified robust associations between TME composition and clinical phenotypes such as tumor subtype and grade that were not discernible when the analysis was limited to individual datasets, highlighting the need for atlas-based analyses. This atlas represents a valuable resource for further high-resolution analyses of TME heterogeneity within BC.

**Graphical Abstract:** 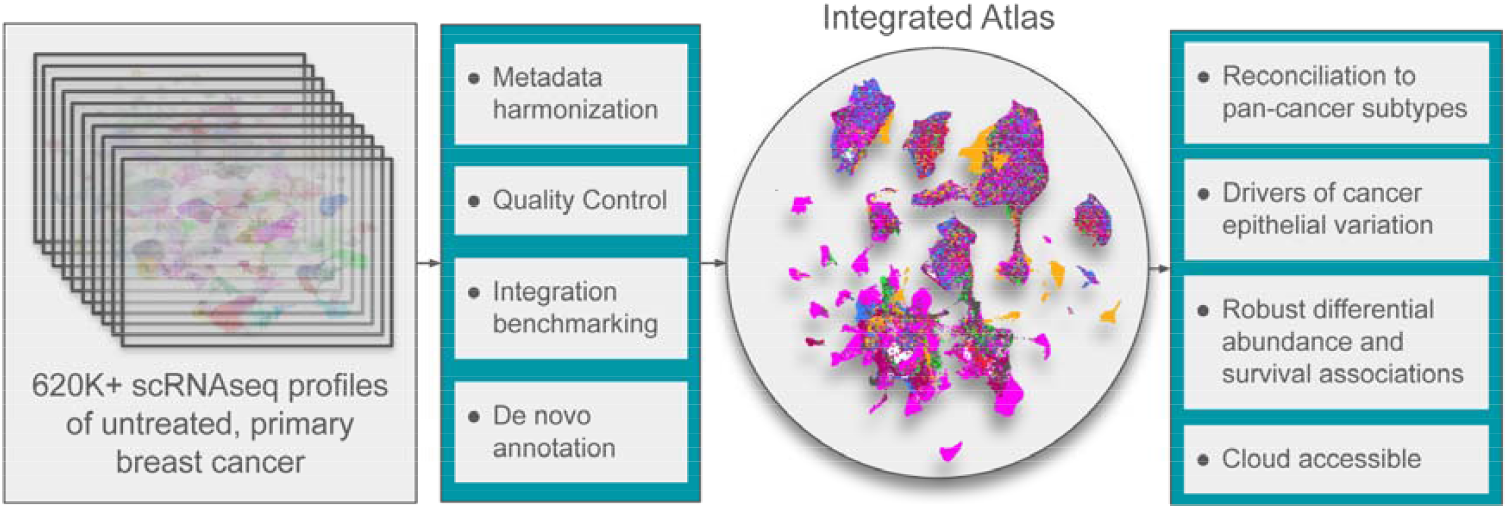

## Introduction

Breast cancer (BC) is the most prevalent cancer and the second most common cause of cancer death in women (1). It is defined as a malignancy of the epithelial duct in breast tissue, and is a highly heterogeneous disease (2, 3) for which clinical outcomes and responsiveness to treatment depend greatly on intrinsic factors related to the cancer cell, as well as extrinsic factors related to altered states of immune and stromal cells in the tumor microenvironment (TME) (4–6). To improve our understanding of breast cancer progression and treatment, an extensive characterization of the heterogeneity in BC cancer cells and the surrounding tumor microenvironment is required.

Single cell transcriptomics (scRNAseq) remains the method of choice to characterize heterogeneity in tissue composition in terms of cell *types* – categories of cells that perform consistent functions, and in terms of cell *states* – categories of transient functions that can be shared across different cell types (7, 8). The technology has been extensively applied to profile healthy breast tissue, as part of the larger human cell atlas effort. There are now highly-resolved scRNAseq atlases that have annotated 700,000-800,000 cells across 55-126 samples yielding new insights into the diversity of immune, stromal, and epithelial populations in healthy breast tissue and how these are linked to clinical metadata (9, 10). However, the application of scRNAseq to profile breast cancer (BC) samples has been limited by cohort sizes, with most studies having fewer than 30 samples (11–20). While these individual studies have laid substantial groundwork in characterizing tumor heterogeneity within BC, their limited sample size poses a challenge with reproducibility and identifying statistically significant associations between tissue composition and clinical metadata.

To address this challenge, we integrated eight publicly available scRNAseq datasets of untreated and unsorted BC samples (11–20) to create an atlas of transcriptional heterogeneity in BC. The analysis of this combined atlas enabled 1) the reconciliation of mislabeled cell types, 2) the identification of novel cell types, and 3) the association of changes in TME heterogeneity with clinical phenotypes such as tumor subtype and grade with increased statistical power (Fig 1a). Another study has attempted to integrate publicly-available scRNAseq BC data to create an atlas but compared to Xu et al. our study only contains unsorted BC samples, allowing for an unbiased characterization of inter-patient heterogeneity, and our atlas has a much a larger sample size (> 600,000 cells), enabling more robust identification of cell types and differentially abundant populations across phenotypes (21).

**Figure 1.**
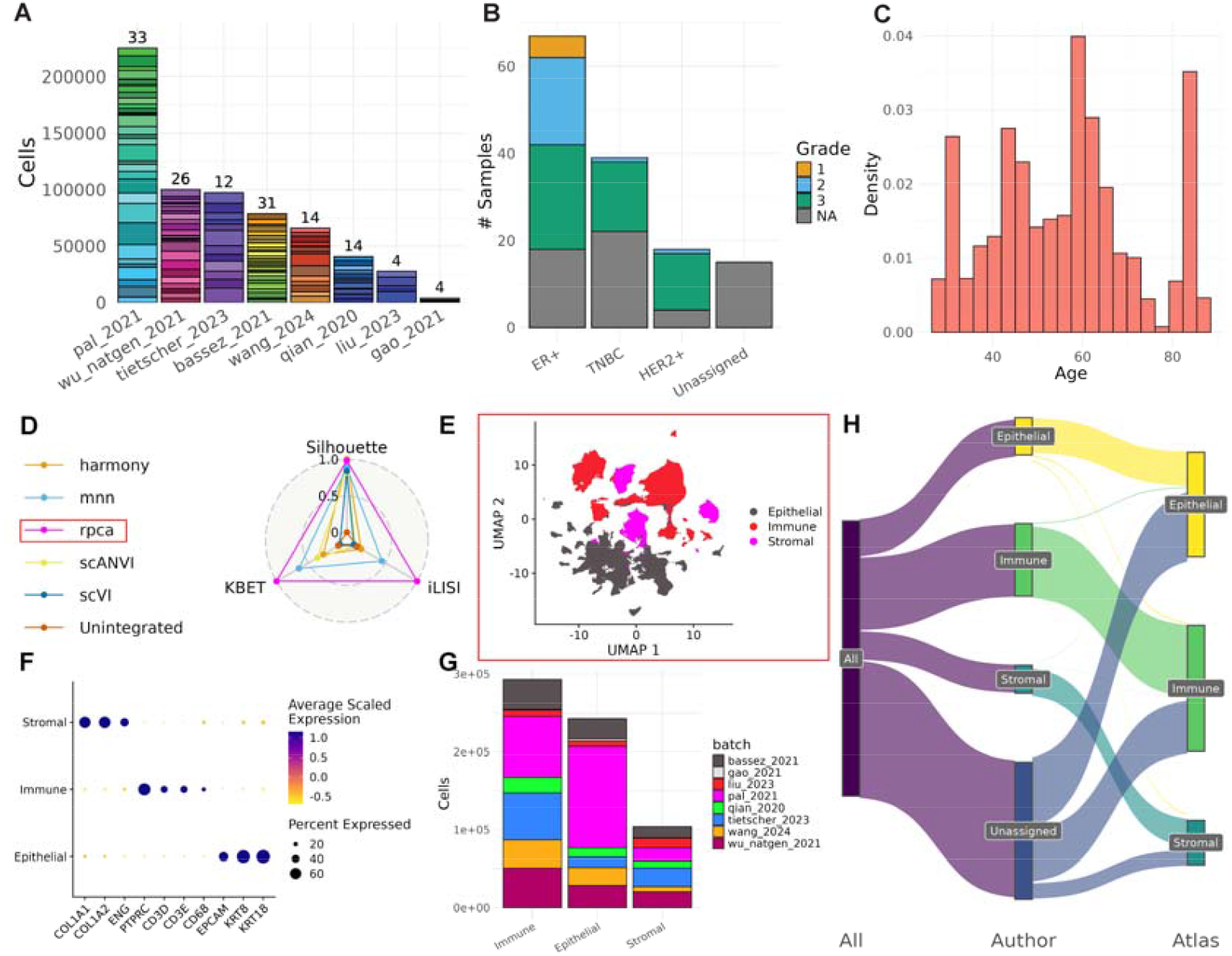
Construction and composition of the integrated breast cancer atlas. A) Distribution of patients and number of cells per patient sequenced across each of the eight datasets. B) Distribution of tumor subtype and grade information. C) Distribution of ages for all patient donors. D) Batch correction metrics for each integration method. E) UMAP of embeddings from chosen integration method (RPCA) and the separation by broad cell type. F) Expression of canonical markers for each of the broad cell types separated from the atlas. G) Distribution of cells from each dataset in each of the three broad compartments. H) Comparison of broad cell type label between author annotations and atlas-based annotations.

This integrated scRNAseq atlas provides an unbiased reference landscape of transcriptional heterogeneity in BC, comprised of more than 600,000 cells across 138 patients, the largest of its kind, and represents a valuable resource for hypothesis-driven analysis of TME heterogeneity. The data and methods described herein provide a robust framework for updating this BC atlas as new datasets are made publicly available (22).

## Methods

### Data Acquisition

To create an unbiased scRNAseq atlas of human BC, we integrated publicly available datasets that contained untreated samples (samples from patients who had not undergone chemotherapy or other treatment before sequencing) and samples that were not sorted prior to sequencing (not biased towards a particular cell type). Given these criteria, we identified eight datasets to include in this atlas, totaling 138 patients and 621,200 cells.

### Data Preprocessing

Each scRNAseq dataset was first pre-processed using Seurat (23) to remove low-quality cells, doublets, and normalize data before integration. We adopted current scRNAseq pre-processing best-practices by using adaptive thresholding based on median absolute deviation, instead of manual cutoffs, to filter out outlier cells based on three criteria: mitochondrial genes expression, number of unique genes detected per cell, and total number of genes detected per cell (24, 25). For studies that only provided pre-filtered data, we retained all cells for downstream analysis. Following filtering, doublet scores were obtained using scDblFinder (26) and expression profiles were normalized by total expression and log-transformed using Seurat’s *LogNormalize* function which has been demonstrated to work well in prior benchmarking (27).

### Integration of Single Cell Data

Integration of filtered scRNAseq data was performed using various methods: Seurat’s V5 reciprocal principal component analysis (RPCA) (23), fastMNN (28), Harmony (29), scVI (30), and scANVI (31). To facilitate fair comparison between the integration methods, the same number of variable features (5000), and PC dimensions (200) were used for integration when possible. For scVI and scANVI, integration is performed at the level of raw counts instead of embeddings, so only the variable feature selection process was included. To avoid biasing the integration towards individual sources of annotations e.g. author annotations or SingleR/CellTypist annotations, only the batch correction metrics: silhouette score, k-nearest neighbor batch effect test (kBET), and integration local inverse Simpson’s index (iLISI), were used to assess integration performance.

### Inference of PAM50 Subtype

To infer molecular subtypes of each sample using the PAM50 method, we first calculated a ‘pseudo-bulk’ profile for each tumor using Seurat’s *AggregateExpression* function, and then applied *molecular*.*subtyping* from the *genefu* R package to each pseudo-bulk profile with default parameters (32). Each pseudobulk profile receives a score normalized between 0 and 1 for each subtype, indicating the likelihood of classification, and the molecular subtype with the highest score leads to the final classification (Supp Table 1A).

**Table 1:** Summary of Datasets used in Atlas.

| Dataset | # Patients | # Cells | System | PMID | Data Link |
| --- | --- | --- | --- | --- | --- |
| Pal et al. 2021 | 33 | 224,823 | 10x 3' | 33950524 | GSE161529 |
| Bassez et al. 2021 | 31 | 78,549 | 10x 5' | 33958794 | Lambrechts lab |
| Wu et al. 2021 | 26 | 99,876 | 10x 3' v2 | 34493872 | GSE176078 |
| Wang et al. 2024 | 14 | 48,553 | 10x 3' v2 | 38382468 | GSE248288 |
| Qian et al. 2020 | 14 | 40,665 | 10x 3' v2 & 10x 5' v2 | 32561858 | Lambrechts lab |
| Tietscher et al. 2023 | 12 | 97,301 | 10x 3' v3 | 36609566 | E-MTAB-10607 |
| Liu et al. 2023 | 4 | 27,490 | 10x 3' v3 | 36594618 | GSE225600 |
| Gao et al. 2021 | 4 | 3,943 | 10x 3' v2 & 10x 5' v2 | 33462507 | GSE148673 |
| **Total** | **138** | **621,200** |  |  |  |

### Discrete annotation of cell types

Annotation of cell types was performed using a combination of reference based annotation methods: singleR (33), CellTypist (34, 35), unsupervised recursive partitioning: K2*Taxonomer* (36), and unsupervised clustering followed by differential expression analysis: MAST (37).

#### Reference-based annotation of cell type

Both singleR and CellTypist require a labelled scRNAseq reference dataset to compare query cells with. Since all of the samples in this study originate from breast tissue, we leveraged labeled scRNAseq data from the healthy breast cell atlas (HBCA) as reference (10). For singleR, the wilcox method *de*.*method* = “*Wilcox*” was used as this is more appropriate than the default methods when applying singleR to single cell data. For CellTypist, we used a pre-trained breast tissue model that used labeled data from HBCA (35), and used majority voting to determine the final cell type label *majority_voting* = *True, mode* = ‘*best match*’.

#### Unsupervised clustering-based annotation

Both K2*Taxonomer* and clustering-based annotation rely on unsupervised clustering which was performed using the Louvain algorithm for three subsets of the atlas separately: epithelial, immune, and stromal, based on expression of canonical markers (Epithelial: *EPCAM, KRT8, KRT 18;* Immune: *PTPRC, CD3D, CD3E, CD68;* Stromal: *COL1A1, COL1A2, ENG*). Performing clustering and annotation within each compartment separately enables the detailed characterization of heterogeneity within each compartment.

To identify differentially expressed genes (DEGs) between clusters, we used Seurat’s *FindAllMarkers* with the parameters *test*.*use = ‘MAST’, latent*.*vars = ‘batch’* to account for the negative-binomial distribution of single cell data and to control for dataset-specific effects respectively.

Whilst clustering-based annotation can quantify differences at the cluster level, the number of clusters, and the resulting DEGs obtained from downstream analysis, relies heavily on arbitrary values of the ‘resolution’ parameter in typical unsupervised clustering algorithms. To mitigate these limitations, we leveraged K2*Taxonomer* to first learn hierarchical relationships between the clusters identified in each compartment, enabling the annotation of clades of clusters instead of just individual clusters. K2*Taxonomer* was applied to the integrated embeddings obtained from RPCA, with the following parameters (*featMetric=“F”, nBoots=400, clustFunc=cKmeansDownsampleSqrt*) which are recommended for large scRNAseq datasets (36). To estimate the differentially expressed genes (DEGs) between nodes in the dendrogram, we used the normalized gene expression matrix and K2Taxonomer’s built in function *runDGEmods*, which uses MAST to model differential expression and the batch variable as a covariate to control for dataset-specific effects.

#### Random Forest-based annotation of discriminative factors

The *randomForest* package was used to train random forests to classify cluster membership, or meta-cluster membership (as determined by K2 *Taxonomer*), of single cell profiles based on expression of continuous scores related to cell stemness, EMP, HBCA cell type, malignancy, and clinical metadata. The model was run with the parameter *importance* = *TRUE* to extract the most important features (as determined by gini-based importances), that differentiated epithelial clusters from each other.

### Continuous annotation of cell states

Classifying discrete cell types is just one facet of heterogeneity, and there are many transient functional programs or cell states that require a more continuous scoring (38). To characterize heterogeneity of functional states within the cancer epithelial compartment of our BC atlas we scored each cell in terms of its malignancy with inferCNV (39), stemness with Cytotrace2 (40), Epithelial-Mesenchymal plasticity (41), expression of hallmark pathways (42, 43), as well as expression of epithelial subtypes identified in HBCA (10).

#### Inference of copy number variations

InferCNV was used to estimate copy number alterations for tumor epithelial cells from each dataset independently with the following parameters: *cutoff* = *0.1, cluster_by_groups* = *TRUE, denoise* = *TRUE* and *HMM* = *FALSE* that are recommended for 10X scRNAseq data. Immune and stromal cells from each dataset were used as normal references, and from these normal reference sets, 100 immune and stromal cells were sampled to act as negative controls to validate inferCNV output.

For each cell, inferCNV estimates a score for each gene that is above one if there is inferred copy number gain, and below one if there is inferred copy number loss. We summarized these scores to estimate a malignancy score per cell that represents the mean absolute CNV score.

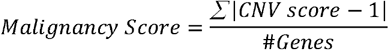

To classify each cell as malignant or non-malignant, for each dataset we identified the 95% quantile of malignancy scores in the non-epithelial cells and labeled all epithelial cells with malignancy scores greater than this cutoff as malignant (Supp Table 1B).

#### Scoring of gene set activity

To characterize the heterogeneity of functional states present in the atlas, we leveraged publicly available transcriptional signatures (Supp Table 1C) and used a rank-based method (AUCell) (44), with default parameters, to score enrichment of signatures in each cell. For the immune compartment, we scored cells using signatures from pan-cancer studies of myeloid and lymphoid cell states (45, 46), as well as signatures of immune cell types from HBCA (10). For the stromal compartment, we scored cells using signatures from a pan-cancer study of stromal cell heterogeneity (47). For the epithelial compartment, we scored cells using canonical hallmark signatures (43), as well as signatures of epithelial cell types from HBCA (10).

### Differential Diversity and Abundance Analysis

#### Cell Type Diversity Analysis

We used an entropy-based cell type diversity metric (CTDS) (48) to assess how the overall diversity in a patient’s TME changes across subtype, age, and tumor grade. The CTDS metric accounts for the number of cells per sample and the compositional nature of proportional data which enables comparison across samples.

#### Differential Abundance Analysis

To test differences in cell type proportions across different phenotype groups e.g. tumor grade and subtype, we used the R package *sccomp* with *bimodal_mean_variability_association* = *TRUE* which is recommended for scRNAseq data (49). To isolate the effects of tumor grade on TME composition, we controlled for tumor subtype, age, and batch with the following formula:

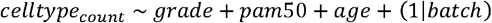

Similarly, to isolate the effects of tumor subtype on TME composition, we controlled for grade, age, and batch.

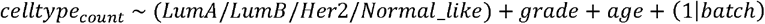

### Multi-resolution survival analysis of cell type populations

To identify the clinical relevance of cell type annotations identified in the atlas, we extracted differentially expressed genes defining each subpopulation found in our taxonomic analysis of each compartment (K2*Taxonomer*) (36), and used the R package *brcasurv* (50) to model the association between the expression of subpopulation specific genes and patient survival in both TCGA (3), METABRIC (51), and SCANB (52).

To address possible confounders in the survival analysis, we included patients age, as well as patient-level gene set projection scores of inflammation and proliferation that have previously been associated with poor prognosis and demonstrated to be necessary for correction in survival modelling (36, 53, 54). The patient-level gene set scores were obtained by using the R package *GSVA* (55) to project inflammation (54), proliferation (53), as well as the cell type specific gene sets onto transcriptional profiles from TCGA and METABRIC. The Cox proportional hazards model underlying the survival analysis controlled for these confounders with the following formula:

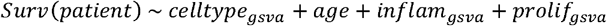

Since the cox proportional hazards model assumes that the hazard ratio is constant over time (proportional hazards assumption), we filtered out results that did not satisfy this criterion (Supp Table 1D). Finally, to control for multiple testing, we corrected p-values in each cox model using the FDR method (56) in the *p*.*adjust* R package, and combined p-values across analyses performed in TCGA, METABRIC, and SCANB using Fisher’s method (57).

## Results

### Constructing the integrated core atlas

The BC atlas was constructed by integrating eight publicly available scRNAseq datasets of untreated primary BC tumors. Across the eight datasets, samples were sequenced at varying throughput levels and had different distributions of clinical and patient metadata including subtype, grade, and age (Fig. 1a,b,c). Prior to integration, batch effects were evident as cells separated by dataset of origin, rather than by biological factors (Supp Fig. 1a). Six different integration approaches were applied and the best-performing method, as determined by atlas integration batch correction metrics (58) was Seurat’s reciprocal PCA method (RPCA) (Fig. 1d).

### Relabeling of broad cell types

To annotate cells in the integrated atlas, we first defined three compartments: epithelial, immune, and stromal, by cross-referencing annotations from SingleR, CellTypist, and author annotations with broad clustering (Louvain clustering with resolution 0.1) (Supp Fig. 1b). The resulting compartments showed distinct expression of canonical marker genes (Epithelial: *EPCAM, KRT8, KRT18;* Immune: *PTPRC, CD3D, CD3E, CD68;* Stromal: *COL1A1, COL1A2, ENG*) and diverse representation across all eight source datasets (Fig. 1e,f,g). More than 300,000 cells that previously had no available cell type annotation were annotated using the cluster-based annotations from the atlas (Fig. 1h).

These annotations of broad cell types based on integrated data also resulted in a relabeling of broad cell types for over 9172 cells (around 1% of the atlas) across 5 datasets (e.g., relabeling a cell as stromal that was originally labeled immune).

### Cancer Epithelial Diversity

Intra-tumor heterogeneity within cancer cells stems from multiple factors including but not limited to genomic instability, epigenetic alterations, and clonal evolution which are all implicated in the treatment of cancer (59). In this BC atlas, we sought to characterize intra and inter-tumor transcriptional heterogeneity by scoring each cancer cell with respect to potency/stemness (40), epithelial-mesenchymal plasticity (EMP) (41), malignancy (39), ‘activity’ of hallmark pathways (43), expression of epithelial cell type markers from HBCA (10), as well as clinical metadata related to age, tumor grade, and subtype (Fig. 2a). Further, we sought to find the driving factors of variation by leveraging a recursive partitioning tree-based method *K2 Taxonomer* (36) to identify meta-clusters and hierarchical relationships between them (Fig. 2b), and also trained random forests to extract the most discriminative features distinguishing each cluster and meta-cluster (Fig. 2c).

**Figure 2.**
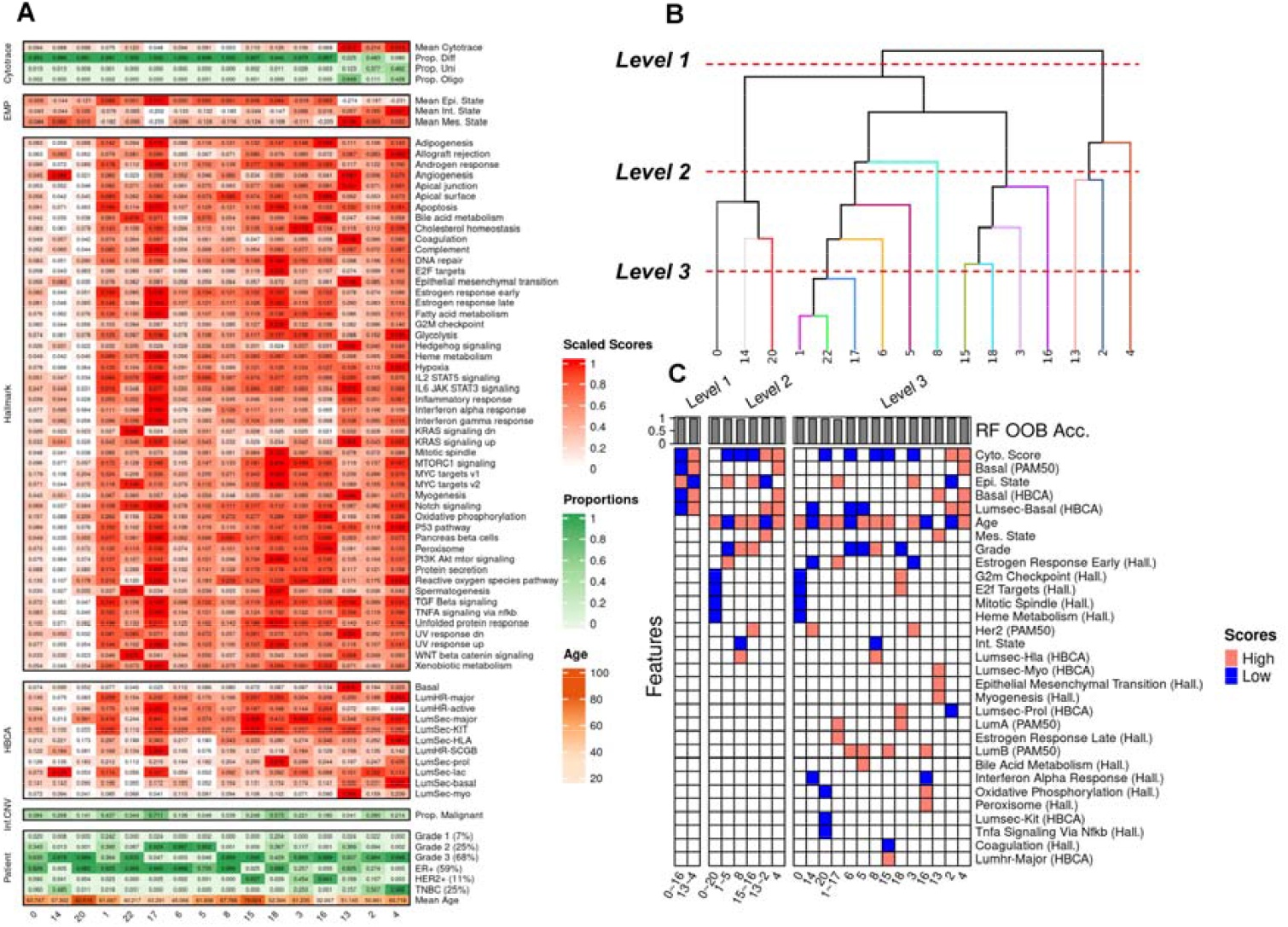
Multi-resolution characterization of Cancer Epithelial Compartment. A) Heatmap of patient metadata, malignancy data (InferCNV), annotation data (HBCA), cell potency data (Cytotrace), plasticity data (EMP), and expression of hallmark pathways (Hallmark). B) Dendrogram of epithelial clusters learned from K2 Taxonomer. C) Top 5 important features, as determined by gini-based importance, from random forest models trained to differentiate different clades within each level of the dendrogram. Red cells indicate that scores related to that feature were higher than the mean in that subset of the dataset.

To identify functional groups of cancer cells shared across patients in the atlas, we performed unsupervised clustering (Louvain clustering with resolution 0.1) of cancer epithelial cells and initially identified 23 clusters that largely stratified by cell typist annotation based on the clinical subtype, and PAM50 inferred subtype (Supp Fig. 2a,b,c). The PAM50 ‘intrinsic’ molecular subtype of a tumor sample is related with, but not equivalent to, clinical annotations of tumor subtype based on immunohistochemistry or in-situ hybridization (Supp Fig. 2d). It provides valuable tumor subtype information especially when clinical subtype data is unavailable.

Though most clusters captured epithelial variation across tumor subtypes and individual donors (Supp Fig. 2e,f), across the range of resolution parameters evaluated (0.1-1.0), even the coarsest resolution of 0.1 yielded multiple clusters (7,9,10,11,12,19,21) mostly consisting of cells from individual patients, as also revealed by their lowest entropy of donor proportions (Supp Fig. 2g,h). This suggests that the variation captured within these clusters was mostly driven by tumor-specific clonal expansion and given our goal of defining shared programs of cancer epithelial heterogeneity, they were excluded from further downstream characterization, leaving sixteen epithelial clusters in the atlas. The differentially expressed genes for each of these sixteen epithelial clusters were identified using *K2 Taxonomer* (36) and MAST (37) (Supp Table 2) and characterized using our taxonomy guided random forest approach described below.

#### Multi-resolution characterization of cancer epithelial cells

Among the remaining 16 epithelial clusters we identified the driving factors of variation within our cancer epithelial compartment by training random forest based classifiers at three separate levels of taxonomic resolution (Fig. 2b,c). In previous integrated single cell studies of breast cancer, cancer epithelial heterogeneity was characterized with respect to PAM50 and clinical subtypes(11, 12), as well as functional annotations by, e.g., Hallmark signatures (21), but has not yet been done in a hierarchical manner to identify the driving factors of variation.

This multi-level characterization revealed segregation of cancer epithelial cells at the highest level (Level 1) by stemness, EMP, and basal phenotypes, with clusters 13 through 4 (13∼4) on the right side of the dendrogram exhibiting more stemness, more EMP (more mesenchymal), and being more basal-like compared to clusters on the left side (clusters 0∼16). At the next level of the dendrogram (Level 2), cancer cells on both sides segregated by age, with the left branch of the tree (clusters 0∼16) segregating by older donors (clusters 0∼20, 8, and 15∼16) and younger donors (clusters 1∼5), and the right branch of the tree (clusters 13∼4) segregating by older (cluster 4) and younger (clusters 13∼2) patients. At the next level of characterization (Level 3), finer groups of cancer epithelial heterogeneity emerge with clusters exhibiting decreased mitotic spindle activity (cluster 0), decreased TNFA signaling (cluster 20), increased EMT activity (cluster 13), as well as subtype-specific clustering (Luminal A: 1∼17, 18; Luminal B: 6, 5, 15, 16; HER2: 3, 14), and grade specific clustering (high grade tumors: 8; low grade tumors: 6, 5, 18).

### Immune and Stromal Cell Diversity in the TME

Across the immune and stromal compartments of the atlas, 31 immune and 14 stromal subpopulations were characterized (Fig. 3a,4a). These subpopulations were identified by integrating data from automatic annotation methods: SingleR (33) and CellTypist (34) (Supp Fig. 3,4), hierarchical tree-based methods (K2 *Taxonomer*) (36) (Fig. 3b,4b), as well as marker genes obtained from differential expression analysis (MAST) (37) (Supp Table 3,4).

**Figure 3.**
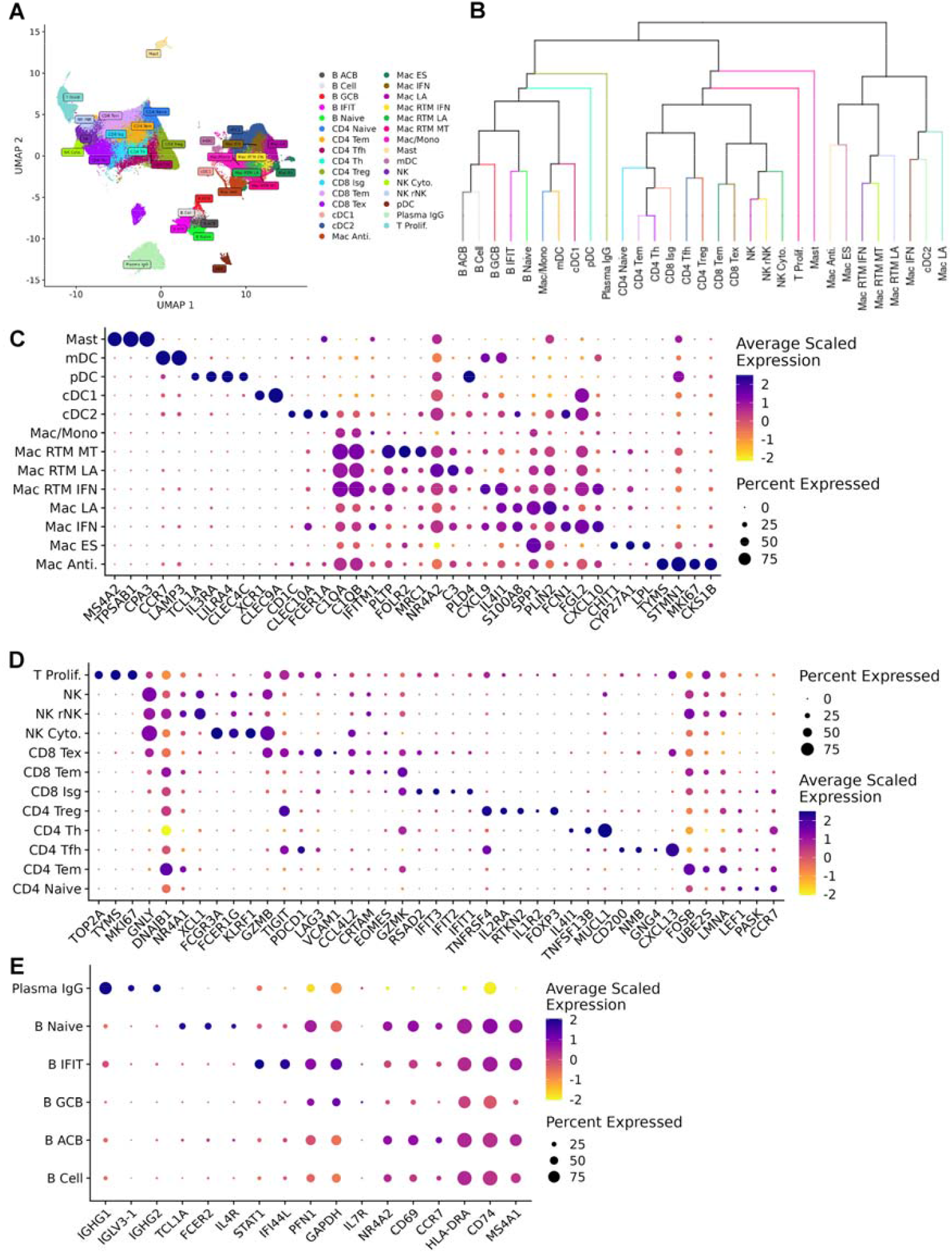
Diversity of immune cell types in the TME. A) UMAP of all immune subtypes characterized in the atlas. B) Dendrogram of immune annotations learned from K2 Taxonomer. C) Dot plot of markers for the myeloid subtypes identified in the atlas. D) Dot plot of markers for the T + NK subtypes identified in the atlas. E) Dot plot of markers for the B + Plasma subtypes identified in the atlas.

**Figure 4.**
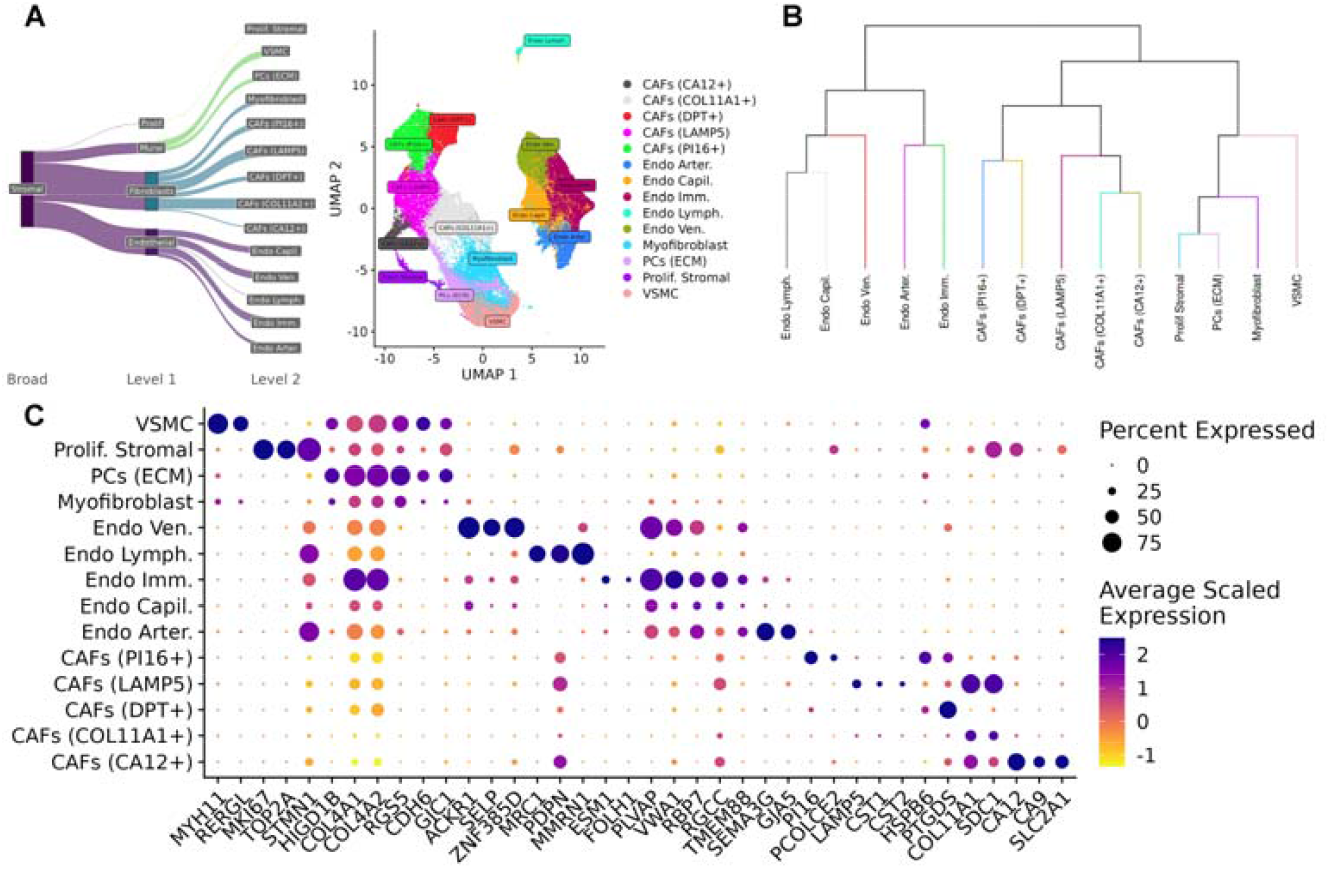
Diversity of stromal cell types in the TME. A) Sankey diagram of all stromal subtypes characterized in the atlas. B) Dendrogram of stromal annotations learned from K2 Taxonomer. C) Dot plot of markers for the stromal subtypes identified in the atlas.

#### Reconciliation of immune cell subtypes to pan-cancer subtypes

Macrophages and monocytes formed eight subpopulations, which were found to have analogs in previous pan-cancer and breast cancer specific characterizations of myeloid cells (10, 45). Using nomenclature from Guimaraes et al. (45), we distinguish between tissue-resident macrophages with *Mac RTM* and circulating monocyte-derived macrophages as *Mac*. In our atlas, we identified two macrophage populations associated with lipid metabolism, *Mac RTM our atlas, we identified two LA* and *Mac LA*, defined by expression of *EGR1, FOSB*, and *SPP1, BNIP3* respectively; two macrophage populations associated with interferon signaling, *Mac RTM* IFN and *Mac IFN*, defined by expression of *CXCL9, CXCL10* and *AREG, FCN1*; a resident tissue macrophage, *RTM* MT, also expressing monocyte markers defined by *GPNMB, LGMN*, and *LIPA;* an early-stage macrophage, *Mac ES*, defined by TIMP3, CHI3L1, ACP5; and an antigen presenting macrophage, *Mac Anti*., defined by *STMN1, HIST1H4C, HMGB2*. The remaining macrophages were not significantly associated with known functional annotations and were thus labeled as general macrophage/monocyte cells *Mac*/*Mono* (Fig. 3a,b,c). While previous studies have characterized heterogeneity of macrophages in terms of FOLR2+ and TREM2+ subtypes (60, 61), our analysis leveraged pan-cancer macrophage signatures to identify eight subpopulations that greatly expand beyond these two known subtypes.

T cells formed nine clusters (Fig. 3a), which were also found to have analogs in previous pan-cancer studies (13, 46, 62). These included CD4 and CD8 effector memory T cells (*CD4/CD8 Tem*), which are characterized by expression of *LMNA, FOS*, and *GZMK, EOMES* respectively; CD4 Tregs defined by expression of *FOXP3, IL2RA*, and *TNFRSF4;* an interferon stimulated group of CD8 T cells (CD8 ISG) defined by strong expression of interferon signaling genes *IFIT1, IFIT2, IFIT3;* naïve CD4 cells characterized by expression of *CCR7, PASK, LEF1;* exhausted CD4 T cells (*CD4 Ex*) defined by expression of canonical exhaustion markers such as *LAG3, HAVCR2, PDCD1* and *TIGIT;* CD4 T follicular helper cells (*CD4 Tfh*) expressing transcription factors *TOX, TOX2* and chemokine *CXCL13;* CD4 T helper cells (*CD4 Th*) which in comparison to *CD4 Tfh* cells expressed high levels of interleukin genes *IL7R*, as well as the chemokine receptor *CCR6*. Finally, a proliferative T cell cluster (*T. Prolif*) was identified exhibiting strong expression of *MIK67* and *STMN1* (Fig. 3,a,b,d).

NK cells formed six clusters (Fig. 3a,d), which were collectively defined by expression of Killer Cell Lectin receptors (*KLRD1, KLRF1*) and growth factor *FGFBP2*. Comparison of these clusters to the NK clusters identified in a previous single cell breast cancer atlas (21) identified two overlapping populations: NK-2 or (NK Cyto), a subpopulation expressing high levels of cytotoxicity-related genes *GZMA, GZMB, PRF1;* and NK-0, or reprogrammed NK cell (*rNK*), which expressed *FOS, JUN, NR4A1*, and *DUSP1*. The reproducibility of these two sub-populations in our larger mega-analysis of single cell breast samples suggests that the cytotoxic and reprogrammed NK cell states are indeed robust sub-populations of NK cells in the breast TME. The remaining four NK subtypes identified in the previous study did not overlap significantly with the markers identified in our atlas (Supp Table 3). We therefore employed a conservative approach to label these remaining clusters as general NK cells.s

#### Mast cells formed one cluster (Fig. 3a,c), defined by expression of the enzymes *TPSB2, CPA3*, and *TPSAB1*

B Cells formed five subpopulations collectively defined by canonical B cell markers such as *CD19, MS4A1, CD74*, as well as expression of human leukocyte antigen genes such as *HLA-DRA* and *HLA-DPB1*. We used the signatures from a previous study characterizing pan-cancer B Cell heterogeneity across 270 patients to identify an activated B Cell population (*B ACB*) defined by expression of *NR4A2, EGR, LY9;* a naïve B cell population (*B Naïve*), defined by expression of *TCL1A, FCER2, IL4R;* a subpopulation expressing interferon-induced genes (*B IFIT*) defined by expression of *IFIT3, ISG15, IFITM1;* and germinal center B cells, defined by expression of *LMO2, SUGCT*, and *ITGB1*. B Cells that were not significantly enriched for functional signatures identified in Ma et al. were labeled as general B cells (Fig. 3,a,e).

#### Identification of dendritic subtypes

Dendritic cells formed four clusters (Fig. 3a,c), which include two conventional dendritic cell populations, one characterized by expression of *CLEC9A* (*cDC1*), and another characterized by expression of *CD1C* and *CLEC10A* (*cDC2*); a mature dendritic cell population (*mDC*) defined by *CCL22, CCR7*, and *CCL19* expression; and a plasMacytoid dendritic cell population (*pDC*) defined by expression of *GZMB, JCHAIN*, and *PTGDS*.

#### Reconciliation of stromal cell subtypes to pan-cancer subtypes

Fibroblasts formed six clusters (Fig. 4a,b,c), which were also reconciled to previously characterized pan-cancer stromal subtypes (47). This includes five clusters of cancer-associated fibroblasts (CAFs) consisting of a population of *COL11A1+ CAFs (COL11A1, COL8A2)* implicated in collagen metabolism (47), a population of *LAMP5 CAFs* defined by expression of *LAMP5* and cystatins *CST1* and *CST2* implicated in promoting EMT (47), a population of PI16+ CAFs defined by expression of PI16 and PCOLCE2, a population of *DPT*+ CAFs defined by expression of *DPT* and *CAPN6*, as well as population of *CA12+* CAFs defined by expression of *CA12* and *SLC2A1* which has been implicated in glycolysis metabolism and hypoxia (47). Apart from these CAFs subtypes, myofibroblasts also formed one cluster by expression of *RGS5* and *CDH6*.

Endothelial cells (ECs) surround blood vessels in the TME which are responsible for angiogenesis and tumor growth and formed a total of five clusters in the atlas (Fig. 5a). Four clusters emerged based on the function of these endothelial cells in supporting vasculature: endothelial vein, arterial, capillary, and lymphatic cells segregated based on expression of different marker genes (Fig. 4a,b,c). A further cluster of immature endothelial cells (*Endo Imm*), defined by expression of *PLVAP, VWA1*, and *CA4*, was identified and has been implicated in poor tumor prognosis (47). Recent pan-cancer studies of stromal cell heterogeneity have implicated SELE+ and CXCR4+ tip cells as the prominent angiogenic and pro-inflammatory endothelial subpopulations in the TME (63). While our endothelial vein cells (*Endo Ven*.) differentially express SELE, we did not identify CXCR4+ endothelial cells in our atlas, perhaps due to the differences between pan-cancer subtypes and breast cancer-specific subtypes.

**Figure 5.**
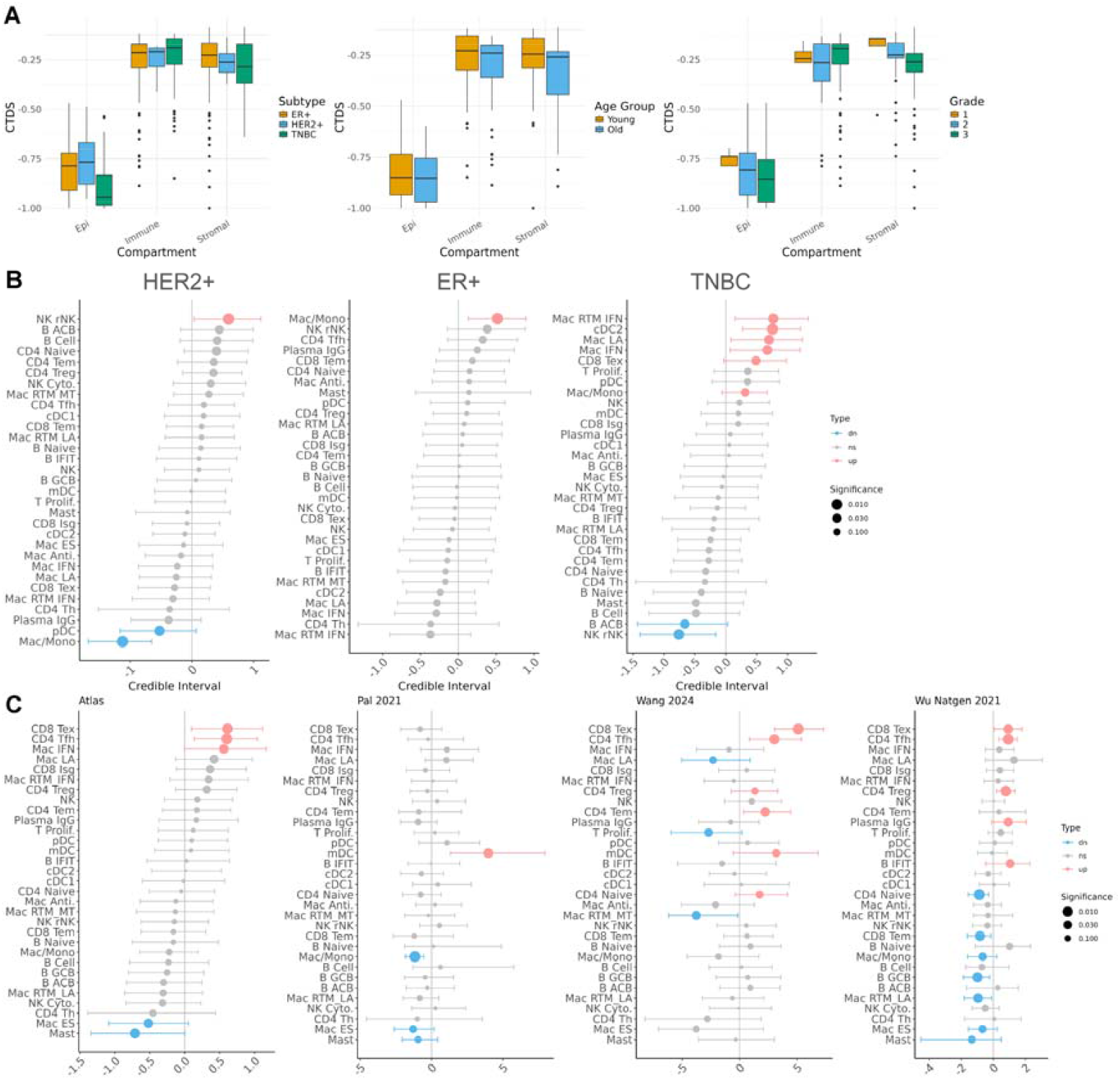
Differential cell type diversity and abundance analysis. A) Cell Type Diversity Score (CTDS) across tumor subtype, patient age, and tumor grade. B) Differential cell type estimates within the immune compartment between HER2+, ER+ and TNBC tumors. C) Differential cell type estimates within the immune compartment between high and low grade tumors.

Other stromal subtypes captured in the atlas include vascular smooth muscle cells (*VSMC*), defined by expression of *RERGL*, and *MYH11;* extracellular matrix related pericytes *(ECM PCs*), defined by expression of collagen genes *COL4A1*, and *COL4A2;* as well as a set of proliferating stromal cells (*Prolif. Stromal*) characterized by high expression of proliferative markers, including *TOP2A, MKI67*.

#### Taxonomic characterization of immune and stromal subtypes

We utilized K2*Taxonomer* to organize the clusters in the immune and stromal compartments into hierarchical taxonomies. Annotating groups of clusters, or clades, mitigates the uncertainty in annotation due to over-clustering. Taxonomic characterization also captures groups of cells sharing similar functions and cell states rather than just cell types.

In the immune compartment, cells largely segregated by lymphoid and myeloid lineages. The exceptions being a population of *dendritic cells*, which clustered with B and plasma cells in a clade due to their shared expression of *antigen presenting genes HLA-DQB1, HLA-DQA1, as well as activation markers CD83* (64), and Mast cells, which clustered within the lymphoid clade due to the lack of expression of macrophage and dendritic markers (*CD68, CD1C*). These exceptions suggest that immune cells in the TME may share similar transcriptional states despite having different lineages (Fig. 3b).

In the stromal compartment, cells partitioned by endothelial and non-endothelial at the first level, and within the non-endothelial clade, further separated between CAFs and mural cells, and lastly between smooth muscle cells, pericytes, and myofibroblasts, indicating strong transcriptome-based partitioning of these stromal subtypes (Fig. 4b).

### Cell type diversity of tumors changes across phenotypes

Using these improved annotations within the atlas, we used an entropy-based cell type diversity score (CTDS) (48) to assess how the overall diversity of the TME changes across patient phenotypes (Fig. 5a). The CTDS score yields the highest value when all cell types (or states) have equal representation in a sample, and the lowest value when a single cell type (or state) is represented in a sample.

Across clinical subtypes, the CTDS of observed epithelial cell states and stromal cell types observed was lowest in TNBC compared to ER+ and HER2+, whereas the average CTDS of immune cell types was highest in TNBC. Using the median age as a cutoff, we compared CTDS across old and young donors and observed a consistent decrease in CTDS across all three compartments in older tumors compared to younger tumors. This expands upon previous studies that identified decreased immune cell type diversity in old subjects compared to young subjects, but only within PBMCs (48). As tumor grade increased, we also observed a consistent decrease in cell type diversity within the epithelial and stromal compartment. The cell type diversity scores provide a global summary of changes in tumor heterogeneity that can complement the more granular differential abundance analysis of individual cell types within the tumor.

### Improved estimates of differential cell type abundance across phenotypes

Using these improved annotations within the atlas, we used *sccomp* (49) to estimate differential cell type abundance across phenotypes such as tumor grade and subtype (Methods).

HER2+ tumors are typically considered to be ‘hot’ tumors with increased immune infiltration compared to other subtypes of tumors (65), but the details of their cell type composition vis-à-vis other tumor subtypes is unclear. In our analysis, we observed that one specific subset of reprogrammed NK cells, *rNK*, increased in abundance in HER2+ tumors relative to other tumor types, whereas *pDCs* and *Mac/Mono* decreased in relative abundance (Fig. 5b). Previous studies have found that pDCs were depleted in HER2+ tumors compared to HER2-tumors (66), and that NK cells were associated with HER2+ tumors (67). However, this is to our knowledge the first report of a specific subpopulation of NK cells found to be associated with HER2+ tumors. With regards to non-immune subpopulations, HER2+ tumors were enriched in immature endothelial cells, and depleted in *CA12*+ and *COL11A1*+ *CAFs* (Supp Fig. 5).

The immune microenvironment of ER+ tumors have been characterized as macrophage-driven (68). In our analysis, we recapitulated this finding and observed that *Mac*/*Mono* were enriched in ER+ tumors compared to other tumors (Fig. 5b). With regards to non-immune subpopulations, ER+ tumors were enriched in *COL11A1*+ *CAFs* and myofibroblasts and were depleted in proliferative stromal cells (Supp Fig. 5).

TNBC tumors are known to exhibit a heterogeneous immune microenvironment, characterized by both increased tumor-infiltrating lymphocytes (TILS) (69), and increased tumor associated macrophages (70). However, specific immune cell types enriched within TNBC have yet to be fully characterized in studies with sufficient statistical power. In our analysis we identified significant enrichment of *Mac RTM IFN, Mac IFN, Mac LA, cDC2*, and *CD8 Tex* cells in TNBC tumors. Conversely, activated B cells and rNK cells were notably depleted. Previous studies have reported expansion of pDCs (71) and depletion of mast cells in TNBC tumors (72). While these trends were also observed in our atlas, their changes were not statistically significant (*fdr* > 0.05) (Fig. 5b). Regarding non-immune cell subpopulations, TNBC tumors were enriched in *CA12*+ *CAFs* and *in Prolif*. Stromal but were depleted in Endo Lymph. (Supp Fig. 5).

Comparison of the atlas-based estimates of differential cell type abundances across tumor grades with analogous estimates derived from the individual datasets further highlighted the utility of the integrated atlas in uncovering unique associations (Fig. 5c). Using our atlas, we not only recapitulated known immune TME associations with tumor grade, such as the enrichment of *CD8 Tex* in higher grade tumors (73), but also identified novel associations, including enrichment of *CD4 Tfh* and *Mac* IFN, alongside depletion of *Mac ES* and *Mast* cells. Beyond confirming previously reported associations, we sought to validate our novel findings using publicly available bulk breast cancer datasets via regression analysis. Of the three bulk breast cancer datasets analyzed in this study (TCGA (3), METABRIC (51), SCANB (52)), only METABRIC had tumor grade data. Within METABRIC, we confirmed positive association between higher tumor grade (grade 3 vs grade 1) and the abundance of *CD8 Tex, CD4 Tfh*, and *Mac IFN*, with both *CD8 Tex* and *Mac IFN* passing the significance threshold (q < 0.05) (Supp Table 1E). For the negative associations identified in our atlas’ grade analysis, we validated the depletion of Mast cells with increasing tumor grade. Although early-stage macrophages (*Mac* ES) were negatively associated with tumor grade in the atlas, they exhibited a positive but non-significant association in METABRIC (p-value > 0.05). Taken together, four out of five of the immune TME associations discovered in our atlas were independently supported by METABRIC data (Supp Table 1E), underscoring the robustness and biological relevance of our integrated approach.

Finally, we found that significant estimates of differential cell type abundance in the atlas were not significant or in discordant directions when tested in individual datasets. For instance, *CD4 Treg* cells were observed to increase with grade in two studies (Wang 2024 and Wu Natgen 2021) but after including all the datasets in the atlas, they did not reach atlas-wide significance. Similarly, *Mast* cells yielded no significant changes in one of the datasets tested (Wang 2024), although when all datasets were combined in the atlas it was the cell type most significantly decreasing with grade. Only three datasets were included in this comparison since datasets need to have both high grade (grade 3) and low-grade tumors (grade 1 & 2) to be included in this differential abundance analysis, which further highlights the need for integration so that diverse patient and clinical metadata can be pooled in mega-analysis-based approaches.

### Multi-resolution cell type associations with patient survival

To assess the clinical relevance of the subpopulations identified in the atlas, we performed survival analyses by projecting bulk RNA expression data from TCGA, METABRIC, and SCANB onto the gene signatures that defined each subgroup within our taxonomic analyses (Fig. 6). To correct for the confounding effects of patient age, proliferation, and inflammation with respect to patient survival (36, 53), the Cox proportional hazard model also included these factors as covariates (Methods).

**Figure 6.**
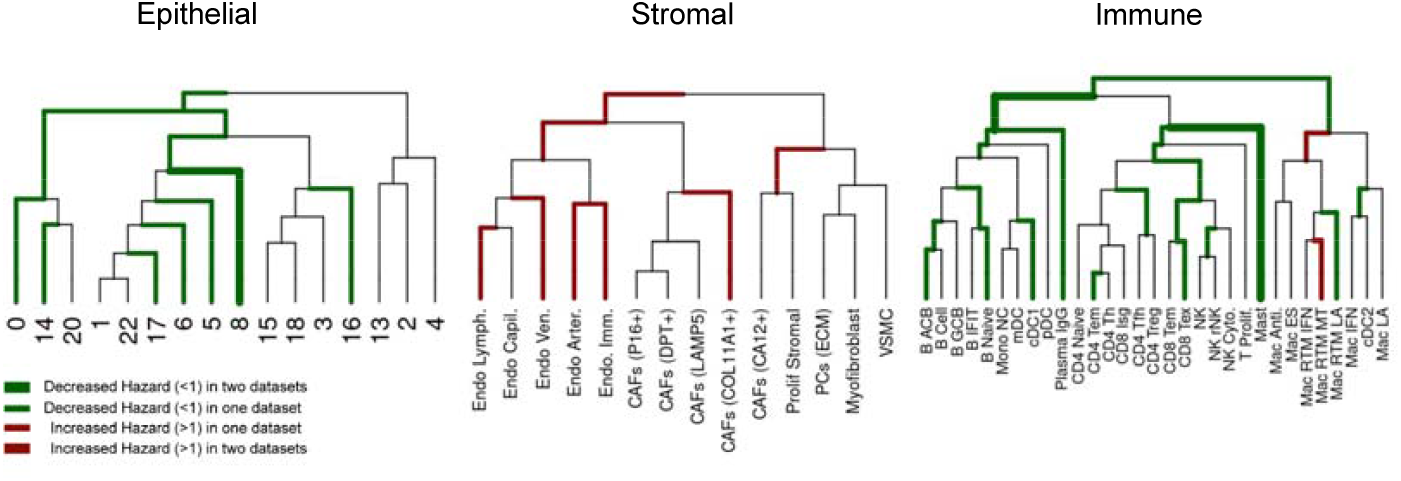
Survival associations of subpopulations found within taxonomic analysis of epithelial, stromal, and immune compartments with respect to TCGA, METABRIC, and SCANB. Green branches indicate a clade that confers a survival advantage (hazard ratio < 1), and red branches indicate a survival disadvantage (hazard ratio > 1) in the respective dataset. The thickness of the branch indicates how many datasets this survival association was statistically significant in.

Two cell populations were significantly associated with survival in at least two of the datasets and both had a positive survival association (hazard ratio less than 1). These include a subpopulation of epithelial cells (Cluster 8) which is characterized by low stemness (more differentiated) and low grade (Fig. 2a,c), and a cluster of *Mast* immune cells (Cluster 1). Both Mast cell infiltration as well as low grade phenotype within tumors have been previously associated with favorable prognosis in BC (74, 75). Though no single cell type/state was associated with decreased survival in at least two datasets separately, when combining the results by fisher’s method, increased CAFs (COL11A1+) activity was collectively associated with decreased survival. Finally, we repeated this analysis within each tumor molecular subtype and identified subtype-specific subpopulations differentially associated with survival (Supp Table 1D).

## Discussion

In this study we integrated scRNAseq data from 138 primary BC tumors collected across eight studies to create the largest transcriptomic atlas of human BC to date. The large number of patients in our integrated atlas, along with the rigorous annotation of both discrete cell types and continuous cell states, will provide a valuable resource for researchers seeking to explore associations of gene expression and or cell type composition with clinical and molecular phenotypes with increased statistical power.

Our characterization of the cancer epithelial compartment of the atlas revealed substantial heterogeneity between cancer cells along the axes of potency/stemness (differentiation continuum), EMP (epithelial-mesenchymal continuum), expression of hallmark and HBCA pathways, as well as clinical phenotypes. By leveraging K2*Taxonomer* and random forest models, we performed a multi-resolution characterization of the cancer epithelial compartment and identified the most discriminative factors differentiating epithelial clusters at different levels of the taxonomy of BC epithelial cells. At the first level, we found that BC tumor cells primarily segregated due to EMP, stemness, and basal phenotypes. At the second level, tumor cells primarily segregated by age, suggesting that aging processes can be a major driver of tumor heterogeneity (76). At the third level, tumor cells separate through more granular hallmark pathways, tumor subtype and grade. While previous studies have employed NMF to identify novel gene programs defining tumor heterogeneity (77), in our study we sought to characterize cancer epithelial heterogeneity using a combination of known epithelial phenotypes i.e. stemness and EMP, clinical metadata i.e. patient age, tumor subtype and grade, and gene programs i.e. hallmarks and HBCA, as such, more flexible methods are needed such as our K2 taxonomy-based random-forest analysis. Further, our multi-resolution characterization of cancer cells has several advantages over typical unsupervised clustering approaches including mitigating the risks of over-clustering and thus over-annotation, as well as being able to identify the major factors driving heterogeneity in a dataset.

In the immune and stromal compartments, our analysis substantially expanded the repertoire of cell type annotations previously reported in individual datasets by integrating insights from pan-cancer studies of myeloid cells, B-lymphocytes, and T cells. Notably, we identified nine distinct T Cell subpopulations, thereby expanding prior characterizations of T cell heterogeneity within the breast TME (78). Additionally, while we reproduced NK cell subpopulations described in an existing breast cancer atlas, our substantially larger sample size enabled us to distinguish which NK subsets are robustly reproducible across cohorts and which may represent dataset-specific observations. Regarding stromal heterogeneity, recent work by Liu et al. utilized spatial transcriptomics to incorporate information from neighboring cells, resolving four spatial CAF subtypes common across cancers (79). In contrast, since our atlas is based solely on transcriptomics data, our annotation strategy relied on pan-cancer stromal signatures, through which we identified six novel CAF subpopulations. This highlights complementary strengths of transcriptomics and spatial approaches for elucidating stromal complexity.

Using the derived annotations of immune cell types and states, we performed cell type diversity and differential abundance analyses to capture global and more granular changes in TME cell type composition with respect to clinical phenotypes such as tumor subtype (ER+, HER2+, TNBC). The cell type diversity analysis revealed an association between increasing tumor grade and decreasing overall TME diversity, as well as age-associated shifts across epithelial, stromal, and immune cells. Differential abundance analysis further resolved immune cell composition differences unique to each subtype, offering insights relevant for therapeutic strategies. The enrichment of *and interferon signaling macrophages* in TNBC suggests promising therapeutic targets, given their respective roles in antigen presentation driving anti-tumor T activation (80) and in shaping a proinflammatory yet immunosuppressive TME (45). The observed expansion of *Mac LA* in TNBC is of interest and may help explain recent findings that TNBC is uniquely responsive to lipid-targeting drugs (81). In HER2+ tumors, the expansion of rNK cells may underlie the efficacy of anti-HER2 therapies known to recruit NK cells (82), underscoring the potential to develop additional NK-targeting treatments, especially since NK cell activity serves as a prognostic biomarker in this subtype. Collectively, these subtype specific differences in the immune TME provide mechanistic insight into the efficacy of existing cancer therapies and may guide prioritization of future therapeutic targets. Importantly, repeating the differential abundance analysis using only individual datasets from the atlas revealed considerable variability across cohorts. This underscores the necessity of data integration to achieve robust and statistically significant associations between cell type composition and clinical phenotypes.

Our atlas-wide survival analysis of cell type subpopulations, utilizing bulk data from TCGA, METABRIC, and SCANB (Supp Table 1D) recapitulated well-established associations, such as decreased mortality risk linked to increased CD4 memory T cell activity and higher proportions of low-grade cancer epithelial cells. Importantly, we also identified novel associations, including increased mortality risk associated with specific endothelial and CAF subpopulations, which may serve as valuable prognostic markers in breast cancer.

A key limitation of this atlas is the absence of adipocytes, despite their recognized importance in cancer progression and aggressiveness (83). This omission stems from technical challenges inherent to scRNAseq, which inadequately captures adipocyte populations. Complementary technologies such as single-nucleus RNA sequencing, and spatial transcriptomics will be essential to characterize adipocyte heterogeneity across patients. Additionally, the atlas represents a static snapshot of the transcriptome and does not capture dynamic changes in response to treatments like radiation or chemotherapy, which could significantly alter the cellular composition and gene expression profiles. Future studies incorporating temporal analyses of cellular responses to various therapies could enhance the atlas’ utility in guiding personalized treatment strategies and understanding mechanisms of treatment resistance.

In summary, this integrated scRNAseq BC atlas, the largest of its kind to date, addresses a critical gap in our understanding of tumor heterogeneity by providing a comprehensive ‘reference’ landscape of cell populations which not only enables statistically powered analyses of cell type diversity, composition, and survival, but also provides a framework for the construction of further atlases to interrogate tumor heterogeneity.

## Data Availability

The atlas is deposited at cellxgene at: https://cellxgene.cziscience.com/collections/9432ae97-4803-4b9f-8f64-2b41e42ad3cb All code is available at: https://github.com/montilab/brca_atlas and figshare https://doi.org/10.6084/m9.figshare.28685012.v1.

## Author contributions

A.C. performed all analyses and wrote the manuscript.

L.K., and S.M. provided computational guidance and feedback. C.S.E., L.K., and G.V.D, provided biological guidance.

S.M. oversaw the project.

The manuscript was edited by all authors.

## Acknowledgments

We thank Gat Rauner, and Yuhan Qiu for feedback regarding the construction and annotation of the atlas. We also thank Corinn Small for providing thorough feedback on making the data accessible on cellxgene. This work was supported by grants from the National Institute of Health (NIH): U01CA243004 (G.V.D. and S.M.), the National Institute of General Medical Sciences of the NIH under award number T32GM100842, and by a gift from Find The Cause Breast Cancer Foundation (84). The content is solely the responsibility of the authors and does not necessarily represent the official views of the NIH.

## References

1. Siegel, R.L., Giaquinto, A.N. and Jemal, A. (2024) Cancer statistics, 2024. CA Cancer J. Clin., 74, 12–49.

2. Perou, C.M., Sørlie, T., Eisen, M.B., et al. (2000) Molecular portraits of human breast tumours. Nature, 406, 747–752.

3. Cancer Genome Atlas Network (2012) Comprehensive molecular portraits of human breast tumours. Nature, 490, 61–70.

4. Polyak, K. (2007) Breast cancer: origins and evolution. J. Clin. Invest., 117, 3155–3163.

5. Lüönd, F., Tiede, S. and Christofori, G. (2021) Breast cancer as an example of tumour heterogeneity and tumour cell plasticity during malignant progression. Br. J. Cancer, 125, 164–175.

6. Place, A.E., Jin Huh, S. and Polyak, K. (2011) The microenvironment in breast cancer progression: biology and implications for treatment. Breast Cancer Res., 13, 227.

7. Kolodziejczyk, A.A., Kim, J.K., Svensson, V., et al. (2015) The technology and biology of single-cell RNA sequencing. Mol. Cell, 58, 610–620.

8. Fleck, J.S., Camp, J.G. and Treutlein, B. (2023) What is a cell type? Science, 381, 733–734.

9. Reed, A.D., Pensa, S., Steif, A., et al. (2024) A single-cell atlas enables mapping of homeostatic cellular shifts in the adult human breast. Nat. Genet., 56, 652–662.

10. Kumar, T., Nee, K., Wei, R., et al. (2023) A spatially resolved single-cell genomic atlas of the adult human breast. Nature, 620, 181–191.

11. Pal, B., Chen, Y., Vaillant, F., et al. (2021) A single-cell RNA expression atlas of normal, preneoplastic and tumorigenic states in the human breast. EMBO J., 40, e107333.

12. Wu, S.Z., Al-Eryani, G., Roden, D.L., et al. (2021) A single-cell and spatially resolved atlas of human breast cancers. Nat. Genet., 53, 1334–1347.

13. Tietscher, S., Wagner, J., Anzeneder, T., et al. (2023) A comprehensive single-cell map of T cell exhaustion-associated immune environments in human breast cancer. Nat. Commun., 14, 98.

14. Bassez, A., Vos, H., Van Dyck, L., et al. (2021) A single-cell map of intratumoral changes during anti-PD1 treatment of patients with breast cancer. Nat. Med., 27, 820–832.

15. Wang, L., Guo, W., Guo, Z., et al. (2024) PD-L1-expressing tumor-associated macrophages are immunostimulatory and associate with good clinical outcome in human breast cancer. Cell Rep. Med., 5, 101420.

16. Qian, J., Olbrecht, S., Boeckx, B., et al. (2020) A pan-cancer blueprint of the heterogeneous tumor microenvironment revealed by single-cell profiling. Cell Res., 30, 745–762.

17. Liu, Y.-M., Ge, J.-Y., Chen, Y.-F., et al. (2023) Combined single-cell and spatial transcriptomics reveal the metabolic evolvement of breast cancer during early dissemination. Adv. Sci. (Weinh.), 10, e2205395.

18. Gao, R., Bai, S., Henderson, Y.C., et al. (2021) Delineating copy number and clonal substructure in human tumors from single-cell transcriptomes. Nat. Biotechnol., 39, 599–608.

19. Wu, S.Z., Roden, D.L., Wang, C., et al. (2020) Stromal cell diversity associated with immune evasion in human triple-negative breast cancer. EMBO J., 39, e104063.

20. Wu, S.Z., Roden, D.L., Al-Eryani, G., et al. (2021) Cryopreservation of human cancers conserves tumour heterogeneity for single-cell multi-omics analysis. Genome Med., 13, 81.

21. Xu, L., Saunders, K., Huang, S.-P., et al. (2024) A comprehensive single-cell breast tumor atlas defines epithelial and immune heterogeneity and interactions predicting anti-PD-1 therapy response. Cell Rep Med.

22. Johnston, K.G., Grieco, S.F., Nie, Q., et al. (2024) Small data methods in omics: the power of one. Nat. Methods, 10.1038/s41592-024-02390-8.

23. Hao, Y., Hao, S., Andersen-Nissen, E., et al. (2021) Integrated analysis of multimodal single-cell data. Cell, 184, 3573-3587.e29.

24. Heumos, L., Schaar, A.C., Lance, C., et al. (2023) Best practices for single-cell analysis across modalities. Nat. Rev. Genet., 24, 550–572.

25. Amezquita, R.A., Lun, A.T.L., Becht, E., et al. (2020) Orchestrating single-cell analysis with Bioconductor. Nat. Methods, 17, 137–145.

26. Germain, P.-L., Lun, A., Garcia Meixide, C., et al. (2021) Doublet identification in single-cell sequencing data using scDblFinder. F1000Res., 10, 979.

27. Ahlmann-Eltze, C. and Huber, W. (2023) Comparison of transformations for single-cell RNA-seq data. Nat. Methods, 20, 665–672.

28. Haghverdi, L., Lun, A.T.L., Morgan, M.D., et al. (2018) Batch effects in single-cell RNA-sequencing data are corrected by matching mutual nearest neighbors. Nat. Biotechnol., 36, 421–427.

29. Korsunsky, I., Millard, N., Fan, J., et al. (2019) Fast, sensitive and accurate integration of single-cell data with Harmony. Nat. Methods, 16, 1289–1296.

30. Lopez, R., Regier, J., Cole, M.B., et al. (2018) Deep generative modeling for single-cell transcriptomics. Nat. Methods, 15, 1053–1058.

31. Xu, C., Lopez, R., Mehlman, E., et al. (2021) Probabilistic harmonization and annotation of single-cell transcriptomics data with deep generative models. Mol. Syst. Biol., 17, e9620.

32. Gendoo, D.M.A., Ratanasirigulchai, N., Schröder, M.S., et al. (2016) Genefu: an R/Bioconductor package for computation of gene expression-based signatures in breast cancer. Bioinformatics, 32, 1097–1099.

33. Aran, D., Looney, A.P., Liu, L., et al. (2019) Reference-based analysis of lung single-cell sequencing reveals a transitional profibrotic macrophage. Nat. Immunol., 20, 163–172.

34. Domínguez Conde, C., Xu, C., Jarvis, L.B., et al. (2022) Cross-tissue immune cell analysis reveals tissue-specific features in humans. Science, 376, eabl5197.

35. Xu, C., Prete, M., Webb, S., et al. (2023) Automatic cell-type harmonization and integration across Human Cell Atlas datasets. Cell, 186, 5876-5891.e20.

36. Reed, E.R. and Monti, S. (2021) Multi-resolution characterization of molecular taxonomies in bulk and single-cell transcriptomics data. Nucleic Acids Res., 49, e98.

37. Finak, G., McDavid, A., Yajima, M., et al. (2015) MAST: a flexible statistical framework for assessing transcriptional changes and characterizing heterogeneity in single-cell RNA sequencing data. Genome Biol., 16, 278.

38. Dance, A. (2024) What is a cell type, really? The quest to categorize life’s myriad forms. Nature, 633, 754–756.

39. Tickle, T., Tirosh, I., Georgescu, C., et al. (2019) inferCNV of the Trinity CTAT Project. https://github.com/broadinstitute/infercnv. Accessed 16 December 2025.

40. Gulati, G.S., Sikandar, S.S., Wesche, D.J., et al. (2020) Single-cell transcriptional diversity is a hallmark of developmental potential. Science, 367, 405–411.

41. Winkler, J., Tan, W., Diadhiou, C.M., et al. (2024) Single-cell analysis of breast cancer metastasis reveals epithelial-mesenchymal plasticity signatures associated with poor outcomes. J. Clin. Invest., 134.

42. Subramanian, A., Tamayo, P., Mootha, V.K., et al. (2005) Gene set enrichment analysis: a knowledge-based approach for interpreting genome-wide expression profiles. Proc. Natl. Acad. Sci. U. S. A., 102, 15545–15550.

43. Liberzon, A., Birger, C., Thorvaldsdóttir, H., et al. (2015) The Molecular Signatures Database (MSigDB) hallmark gene set collection. Cell Syst, 1, 417–425.

44. Aibar, S., González-Blas, C.B., Moerman, T., et al. (2017) SCENIC: single-cell regulatory network inference and clustering. Nat. Methods, 14, 1083–1086.

45. Guimarães, G.R., Maklouf, G.R., Teixeira, C.E., et al. (2024) Single-cell resolution characterization of myeloid-derived cell states with implication in cancer outcome. Nat. Commun., 15, 5694.

46. Chu, Y., Dai, E., Li, Y., et al. (2023) Pan-cancer T cell atlas links a cellular stress response state to immunotherapy resistance. Nat Med, 29, 1550–1562.

47. Du, Y., Shi, J., Wang, J., et al. (2024) Integration of Pan-Cancer Single-Cell and Spatial Transcriptomics Reveals Stromal Cell Features and Therapeutic Targets in Tumor Microenvironment. Cancer Res., 84, 192–210.

48. Karagiannis, T.T., Monti, S. and Sebastiani, P. (2022) Cell Type Diversity Statistic: An Entropy-Based Metric to Compare Overall Cell Type Composition Across Samples. Front. Genet., 13, 855076.

49. Mangiola, S., Roth-Schulze, A.J., Trussart, M., et al. (2023) sccomp: Robust differential composition and variability analysis for single-cell data. Proc. Natl. Acad. Sci. U. S. A., 120, e2203828120.

50. Chen, A. (2025) brcasurv. https://github.com/montilab/brcasurv. Accessed 16 December 2025.

51. Curtis, C., Shah, S.P., Chin, S.-F., et al. (2012) The genomic and transcriptomic architecture of 2,000 breast tumours reveals novel subgroups. Nature, 486, 346–352.

52. Staaf, J., Häkkinen, J., Hegardt, C., et al. (2022) RNA sequencing-based single sample predictors of molecular subtype and risk of recurrence for clinical assessment of early-stage breast cancer. NPJ Breast Cancer, 8, 94.

53. Venet, D., Dumont, J.E. and Detours, V. (2011) Most random gene expression signatures are significantly associated with breast cancer outcome. PLoS Comput. Biol., 7, e1002240.

54. Winslow, S., Leandersson, K., Edsjö, A., et al. (2015) Prognostic stromal gene signatures in breast cancer. Breast Cancer Res., 17, 23.

55. Hänzelmann, S., Castelo, R. and Guinney, J. (2013) GSVA: gene set variation analysis for microarray and RNA-seq data. BMC Bioinformatics, 14, 7.

56. Benjamini, Y. and Hochberg, Y. (1995) Controlling the false discovery rate: a practical and powerful approach to multiple testing. Journal of the royal statistical society series b-methodological, 57, 289–300.

57. Fisher, R.A. (1938) Statistical methods for research workers 7th ed., rev.enl. Oliver and Boyd, Edinburgh.

58. Luecken, M.D., Büttner, M., Chaichoompu, K., et al. (2022) Benchmarking atlas-level data integration in single-cell genomics. Nat. Methods, 19, 41–50.

59. Sun, X.-X. and Yu, Q. (2015) Intra-tumor heterogeneity of cancer cells and its implications for cancer treatment. Acta Pharmacol. Sin., 36, 1219–1227.

60. Komohara, Y., Kurotaki, D., Tsukamoto, H., et al. (2023) Involvement of protumor macrophages in breast cancer progression and characterization of macrophage phenotypes. Cancer Sci., 114, 2220–2229.

61. Wu, S., Jiang, B., Li, Z., et al. (2025) Unveiling the key mechanisms of FOLR2+ macrophage-mediated antitumor immunity in breast cancer using integrated single-cell RNA sequencing and bulk RNA sequencing. Breast Cancer Res., 27, 31.

62. Zheng, L., Qin, S., Si, W., et al. (2021) Pan-cancer single-cell landscape of tumor-infiltrating T cells. Science, 374, abe6474.

63. Li, J., Wang, D., Tang, F., et al. (2024) Pan-cancer integrative analyses dissect the remodeling of endothelial cells in human cancers. Natl. Sci. Rev., 11, nwae231.

64. Hock, B.D., Kato, M., McKenzie, J.L., et al. (2001) A soluble form of CD83 is released from activated dendritic cells and B lymphocytes, and is detectable in normal human sera. Int. Immunol., 13, 959–967.

65. Perrone, M., Talarico, G., Chiodoni, C., et al. (2021) Impact of immune cell heterogeneity on HER2+ breast cancer prognosis and response to therapy. Cancers (Basel), 13, 6352.

66. Łazarczyk, A., Streb, J., Glajcar, A., et al. (2023) Dendritic cell subpopulations are associated with prognostic characteristics of breast cancer after neoadjuvant chemotherapy-an observational study. Int. J. Mol. Sci., 24, 15817.

67. Bouzidi, L., Triki, H., Charfi, S., et al. (2021) Prognostic value of natural killer cells besides tumorinfiltrating lymphocytes in breast cancer tissues. Clin. Breast Cancer, 21, e738–e747.

68. Onkar, S., Cui, J., Zou, J., et al. (2023) Immune landscape in invasive ductal and lobular breast cancer reveals a divergent macrophage-driven microenvironment. Nat. Cancer, 4, 516–534.

69. Fan, Y. and He, S. (2022) The characteristics of tumor microenvironment in triple negative breast cancer. Cancer Manag. Res., 14, 1–17.

70. Yuan, Z.-Y., Luo, R.-Z., Peng, R.-J., et al. (2014) High infiltration of tumor-associated macrophages in triple-negative breast cancer is associated with a higher risk of distant metastasis. Onco. Targets. Ther., 7, 1475–1480.

71. Oshi, M., Newman, S., Tokumaru, Y., et al. (2020) Plasmacytoid dendritic cell (pDC) infiltration correlate with tumor infiltrating lymphocytes, cancer immunity, and better survival in triple negative breast cancer (TNBC) more strongly than conventional dendritic cell (cDC). Cancers (Basel), 12, 3342.

72. Majorini, M.T., Colombo, M.P. and Lecis, D. (2022) Few, but efficient: The role of mast cells in breast cancer and other solid tumors. Cancer Res., 82, 1439–1447.

73. Egelston, C.A., Guo, W., Tan, J., et al. (2022) Tumor-infiltrating exhausted CD8+ T cells dictate reduced survival in premenopausal estrogen receptor-positive breast cancer. JCI Insight, 7.

74. Aponte-López, A., Fuentes-Pananá, E.M., Cortes-Muñoz, D., et al. (2018) Mast cell, the neglected member of the tumor microenvironment: Role in breast cancer. J. Immunol. Res., 2018, 2584243.

75. Jin, Y., Tan, A., Feng, J., et al. (2021) Prognostic impact of memory CD8(+) T cells on immunotherapy in human cancers: A systematic review and meta-analysis. Front. Oncol., 11, 698076.

76. Chatsirisupachai, K., Lagger, C. and de Magalhães, J.P. (2022) Age-associated differences in the cancer molecular landscape. Trends Cancer, 8, 962–971.

77. Gavish, A., Tyler, M., Simkin, D., et al. (2021) The transcriptional hallmarks of intra-tumor heterogeneity across a thousand tumors. bioRxiv, 10.1101/2021.12.19.473368.

78. Azizi, E., Carr, A.J., Plitas, G., et al. (2018) Single-Cell Map of Diverse Immune Phenotypes in the Breast Tumor Microenvironment. Cell, 174, 1293-1308.e36.

79. Liu, Y., Sinjab, A., Min, J., et al. (2025) Conserved spatial subtypes and cellular neighborhoods of cancer-associated fibroblasts revealed by single-cell spatial multi-omics. Cancer Cell, 43, 905-924.e6.

80. Sebastião, A.I., Simões, G., Oliveira, F., et al. (2025) Dendritic cells in triple-negative breast cancer: From pathophysiology to therapeutic applications. Cancer Treat. Rev., 133, 102884.

81. Scully, T., Kase, N., Gallagher, E.J., et al. (2021) Regulation of low-density lipoprotein receptor expression in triple negative breast cancer by EGFR-MAPK signaling. Sci. Rep., 11, 17927.

82. Muntasell, A., Rojo, F., Servitja, S., et al. (2019) NK cell infiltrates and HLA class I expression in primary HER2+ breast cancer predict and uncouple pathological response and disease-free survival. Clin. Cancer Res., 25, 1535–1545.

83. Gao, Y., Chen, X., He, Q., et al. (2020) Adipocytes promote breast tumorigenesis through TAZ-dependent secretion of Resistin. Proc. Natl. Acad. Sci. U. S. A., 117, 33295–33304.

84. Find The Cause Breast Cancer Foundation (2016). https://findthecausebcf.org/. Accessed 16 December 2025.

